# A wildlife behaviour detection model: a case study on the red-legged seriema

**DOI:** 10.64898/2026.08.02.742331

**Authors:** Lotte Schroth, Filipe R.C. Cunha

## Abstract

Object detection models based on deep learning are increasingly used to reduce the manual workload associated with the processing of video recordings in behavioural research, but reproducible workflows describing how ecologists can build such models themselves remain scarce. Here, we present a step-by-step workflow for constructing a species-specific detection model and apply it as a case study to the red-legged seriema (*Cariama cristata*), a widely distributed but poorly studied Neotropical bird. Using 60 videos recorded with camera traps as well as non-stop recording cameras at the campus of the Universidade Federal de Viçosa (Florestal, Minas Gerais, Brazil) between 2023 and 2026, we extracted and annotated 480 images to train a YOLO26-nano detection model. On the validation set, the model achieved a precision of 0.934 and a recall of 0.786 (mAP@50 = 0.870), while performance on the independent test set was slightly lower, with a precision of 0.865 and a recall of 0.740 (mAP@50 = 0.740). When applied to unseen footage, the model processed video at roughly 4.6 times real-time speed on a standard CPU and correctly classified all four test videos for the presence or absence of seriemas. Most missed detections involved seriemas that were distant, partially out of frame, or recorded with the lower-quality non-stop cameras. These results showed that a usable detection model can be built from a comparatively small curated dataset, and the accompanying workflow and code are intended to lower the barrier for ecologists seeking to apply object detection to their own study species.

## Introduction

The use of video recordings has become increasingly common in biological research, especially in conservation and behavioural studies (Caravaggi et al., 2017; Rovero et al., 2013). The use of camera traps is particularly common in this line of research; these typically rely on a passive infrared sensor that triggers the start of a recording when movement is detected (Apps & McNutt, 2018). A well-known issue with camera traps in the wild, however, is that this trigger is not always reliable: false triggers lead to videos or images without any animal present, which drains the camera’s battery (Apps & McNutt, 2018) and can cost hours of post-hoc analysis. Conversely, the trigger may fail to activate or have a delayed activation, causing valuable observations to be missed partially or entirely (Apps & McNutt, 2018). Where logistically possible, researchers sometimes chose non-stop video surveillance cameras instead of camera traps, as these carry less risk of animals going unrecorded (Pechacek, 2005; Reif & Tornberg, 2006). Research comparing the performance of camera traps to non-stop recording cameras found that camera traps missed 43.6% of small mammals and 17% of medium-sized mammals, indicating that camera traps can indeed miss a substantial amount of data (Jumeau et al., 2017). Non-stop recording cameras, however, require a continuous power supply, as they use considerably more energy, and the volume of footage they generate is large, making post-field video analysis extremely laborious. It is still common for footage from wildlife cameras to be sorted by hand, which is inefficient and time-consuming. Recently, deep learning models have attracted growing interest as a way of automating object detection and streamlining footage selection for wildlife research (Norouzzadeh et al., 2018; Shepley et al., 2021; Tan et al., 2022). Modern object detection frameworks, such as YOLO (Redmon et al., 2016), FCOS (Tian et al., 2019) or MMDetection (Chen et al., 2019), can accurately detect and distinguish between animals while processing large quantities of data. These models are trained on large amounts of data on the species of interest, allowing them to learn to recognise visual patterns, and can subsequently detect both the location and class of the objects on which they were trained (Zhao et al., 2019). These developments offer an opportunity to reduce the amount of manual video annotation and labour required in behavioural and conservation research.

Despite these models becoming more common in behavioural ecology, implementing an object detection model remains challenging for many ecologists. Moreover, most studies on animal detection models present only species-specific results, without the description of a reproducible workflow. There is also a taxonomic bias in biodiversity data, with many species being poorly represented (Cunha et al., 2023). As deep learning models generally require large, well-annotated datasets for training, developing accurate detection models for rare and understudied species remains challenging (Villon et al., 2022).

Although animal detection models have been developed and used for several wildlife species, practical workflows that describe how researchers can construct such a reliable model themselves remain lacking. This study addresses this gap by developing and evaluating a YOLO26-based detection model for red-legged seriemas (*Cariama cristata*), a poorly studied Neotropical bird species, while providing a reproducible workflow that can be adapted to other wild species. The red-legged seriema is a large bodied terrestrial bird, widely distributed across South America, including Argentina, Paraguay, Bolivia and Brazil (Jones et al., 2024). These birds inhabit open Cerrado and wetland areas (Brooks, 2014). Although considered a common species information about their ecology remains largely unknown, especially when it comes to behavioural traits and territoriality. Continuous video monitoring provides a tool to better analyse the behaviour of seriemas, but large volumes of data would still require efficient processing. A reliable seriema-detection model aids the efficiency of video selection and thus facilitates research towards this species. This study aimed at providing a step-by-step, reproducible workflow for creating an animal detection model, we then furtherevaluated this pipeline on a poorly studied species as a proof of concept.

## Methods

### Study area and data organization

For the construction of the seriema-detection model, this study used video recordings collected between 2023 and 2025 on the campus of the Universidade Federal de Viçosa (UFV) in Florestal, Minas Gerais, Brazil. These videos were recorded with Bushnell camera traps, producing recordings between 30 seconds and 1 minute in length, saved at a resolution of 1920 × 1080 pixels and a frame rate of 30 fps. Additional footage was obtained at UFV using Green Feathers solar-powered Wi-Fi non-stop cameras between 2025 and 2026. These non-stop cameras produced videos at a resolution of 2304 × 1296 pixels and a frame rate of 11 fps. Although the Green Feathers recordings had a higher nominal resolution than the Bushnell videos, visual inspection indicated a lower apparent video quality, with more visible image noise.

The non-stop cameras saved footage as media files onto a local SD card; these were later extracted as MP4 files using the software ‘crittervid, which split the footage into 50-minute-long clips (Zukas, n.d.). Both the camera-trap and non-stop videos were recorded at multiple locations across the campus, with at least two recordings per location included in this study.

### Dataset construction

In total, 60 videos were used for this study, consisting of 13 non-stop camera videos and 47 camera trap videos. The videos were split into a training, validation and test set, to prevent data leakage. As the dataset was relatively small, the target split ratio was set to 60% training, 20% validation and 20% test. Stratification was used to evenly distribute videos with and without seriemas present, and across the different recording locations.

For each video, the year, location, and presence or absence of seriemas were recorded. Using R (R Core Team, 2026) and RStudio (Posit team, 2026), we used the function createDataPartition from the ‘caret’ package (Kuhn et al., 2024) to stratify the data splits, with ‘set. seed(123)’, used to ensure reproducibility. Because of the small sample size, the resulting split deviated from the target 60/20/20 proportions, initially yielding 40 videos for training, 12 for validation and eight for testing. A small number of videos were therefore manually reassigned between sets to achieve the intended proportions, while maintaining stratification for location and seriema presence. This resulted in 36 videos in the training set and 12 videos in both the validation and test set.

The training set consisted of 30 camera trap videos and six non-stop camera videos. The validation set contained eight camera trap videos and four non-stop camera videos, while the test set contained nine camera trap videos and three non-stop camera recordings.

### Extracting frames from videos

To train the model, images first needed to be extracted from the selected videos. This was done with a custom Python script (available in the associated repository) based on the OpenCV library (Bradski, 2000). For each video, eight frames were selected, evenly distributed across the length of the video. These frames were added to a new folder labelled ‘images’, to keep them easily accessible for the next step.

### Labelling images

For this study we used a Python environment with version 3.14.6 (Python Software Foundation, 2025). The model construction process relied on several packages, most importantly Labelme (Russell et al., 2008), Ultralytics YOLO (Ultralytics, 2026) and OpenCV (Apache Software License, 2026).

Once selected, the images had to be annotated according to the presence of the species of interest — in this study, the red-legged seriema. This was performed using LabelMe, where all images were checked, and bounding boxes were drawn as closely as possible around any seriemas present. If it was difficult to determine from an image whether something was an actual seriema, no bounding box was drawn. Where images contained multiple seriemas, multiple bounding boxes were drawn (Figure 1). This process produced a corresponding .json file for every image containing seriemas, saved directly in the images folder.

**Figure 1.**
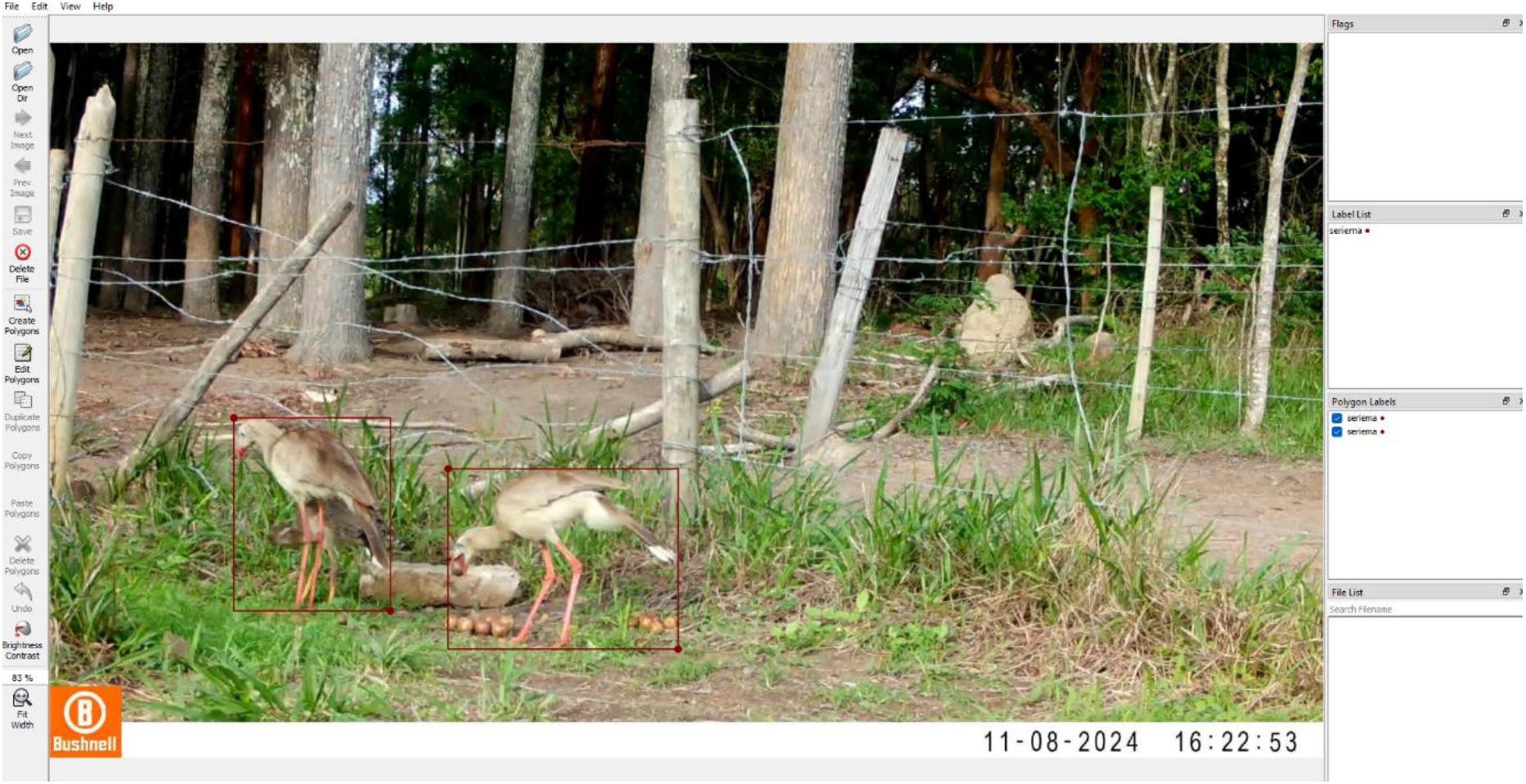
Example of a bounding box drawn around a red-legged seriema during image annotation in LabelMe.

Detection models should not only be trained on the species of interest but also on other animals or structures that may be present in the study area. For this reason, in addition to images containing seriemas, the dataset also included empty images and images of other species present in the area, such as the crested caracara (*Caracara plancus*) and the South American coati (*Nasua nasua*). No bounding boxes were drawn for these images, as detecting these species was not the aim of this study; however, the model still requires a corresponding, empty .json file for each of these images to function correctly.

These empty .json files were created with another custom Python script (available in the associated repository) and saved in the same folder as the images and the .json files containing bounding boxes, allowing a straightforward check that every frame had a corresponding .json file.

### Converting JSON annotations to YOLO format

Given that YOLO models (Jocher et al., 2026) cannot process .json files directly, these first needed to be converted into YOLO-format label files. Scripts provided by Ultralytics are available for this purpose, rely on generating an additional data split. As the dataset had already been split, we converted the .json files into YOLO-format files using a Python script, which is available in the associated repository.

### Creating the YAML configuration file

Finally, we created a YAML file to inform the model where the training, validation, and test sets were stored, and to assign names to the different category IDs. As this model was only intended to identify seriemas, a single category ID (‘0’) was used and given the corresponding name.

### Training the YOLO model

This study used a YOLO model — ‘You Only Look Once’ which can be trained relatively quick, as they are pre-trained on ImageNet (Jiang et al., 2022). We used YOLO26 (Jocher et al., 2026), the most recent YOLO version at the time of this study, specifically the nano variant, which is the fastest but least accurate configuration of YOLO26.

The model was trained for 100 epochs, with a batch size of eight and an image resolution of 640 pixels. No custom data augmentation was used, meaning the default YOLO26 settings were applied. During inference, the default setting of Non-Maximum Suppression (NMS) of 0.7 was retained. Once these settings were defined, the model could be trained; training time is varied and is dependenton the hardware used and on whether a graphics processing unit (GPU) or a central processing unit (CPU) is used. For this study, a CPU was used, which increased training time, but did not affect the outcome of the model. The training procedure uses only the training and validation sets; the resulting model can then be evaluated on the held-out test set to assess its performance.

### Applying the model to new footage

The trained model can also be applied to full videos rather than individual images. For longer videos, it can be useful to avoid scanning through every single frame; the Python script used here (available in the associated repository) therefore applied a video stride of 10 frames, which speeds up processing considerably.

For this study, the model was applied to four videos to assess the average inference time per frame, the total processing time per video, and the overall performance in identifying seriemas.

## Results

Using LabelMe, bounding boxes were drawn on all images in which red-legged seriemas were present. With the trained detection model, predictions could then be made on the presence of seriemas in new images. Model performance was assessed by examining precision as well as the mean average precision at an intersection-over-union (IoU) threshold of 0.5 and across an IoU threshold of 0.5-0.95. These metrics were calculated across confidence thresholds to evaluate overall model performance on both the validation and test sets. All performances were slightly higher on the validation set than for the test set (Table 1).

**Table 1:**
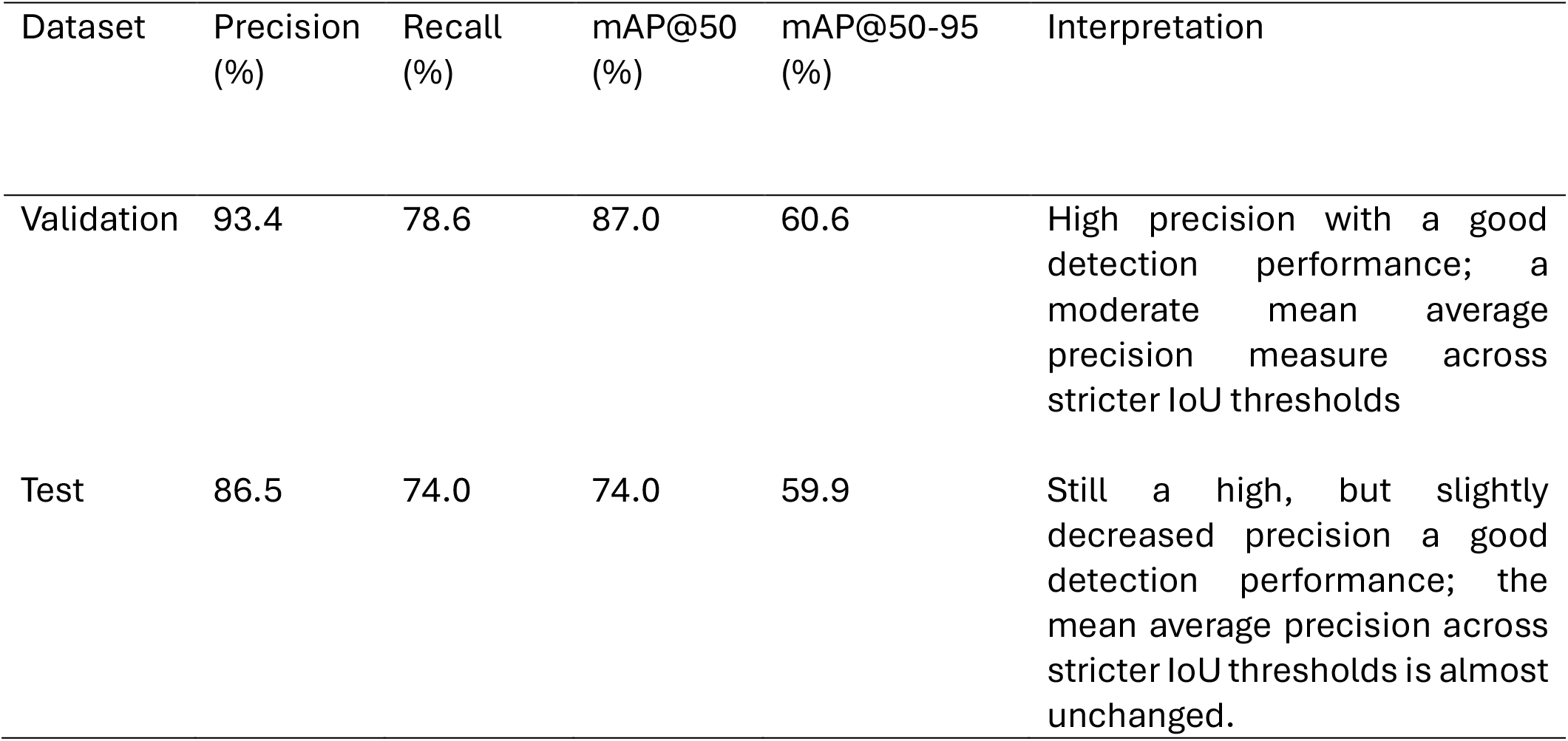
Performance of the YOLO26n model across confidence intervals for detecting red-legged seriemas on both the validation and independent test set. Precision and recall describe the detection accuracy, while mAP@50 and mAP@50-95 describe the overall detection and localisation performance.

| Dataset | Precision (%) | Recall (%) | mAP@50 (%) | mAP@50-95 (%) | Interpretation |
| --- | --- | --- | --- | --- | --- |
| Validation | 93.4 | 78.6 | 87.0 | 60.6 | High precision with a good detection performance; a moderate mean average precision measure across stricter IoU thresholds |
| Test | 86.5 | 74.0 | 74.0 | 59.9 | Still a high, but slightly decreased precision a good detection performance; the mean average precision across stricter IoU thresholds is almost unchanged. |

The validation set consisted of 96 images, containing 56 annotated seriemas in total. At a confidence threshold of 25%, the model was able to correctly identify 45 seriemas, meaning 11 seriemas were missed, resulting in false negatives (Figure 2). The model also produced nine false positives, where other structures were identified as seriemas, most of which involved other species being mistakenly annotated as seriemas, such as dogs and coatis.

**Figure 2.**
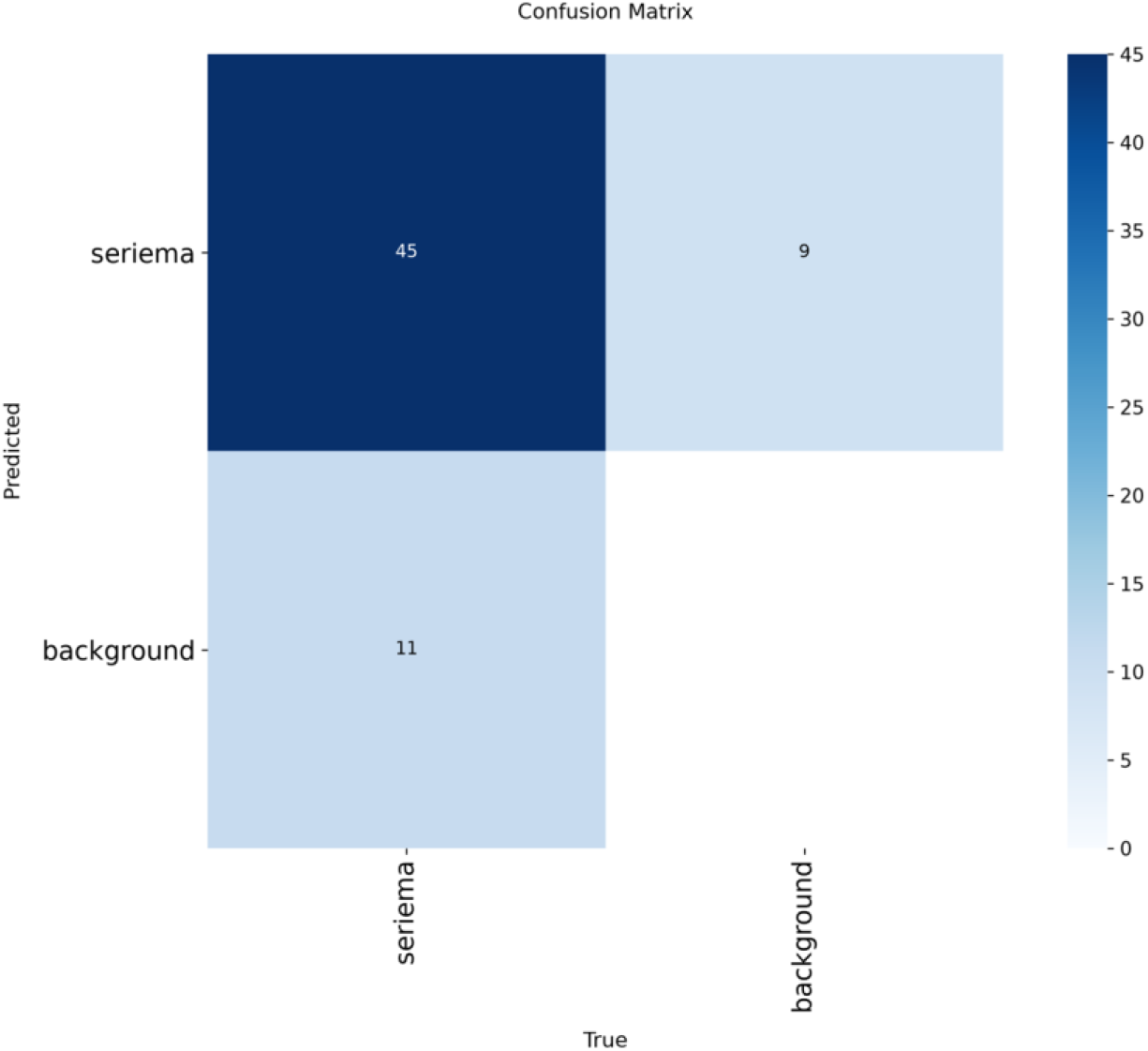
Confusion matrix at a confidence threshold of 25% showing model predictions for the categories seriema and background in the validation set, including true positives, false positives, true negatives and false negatives.

The test set again consisted of 96 images and contained 78 annotated seriemas. At a confidence threshold of 25%, the model correctly identified 56 of these instances, with 22 false negatives and 4 false positives (Figure 3). Of the false positives, only one involved an actual different animal; the other three cases were seriemas that had not been fully annotated in the original dataset. The false negatives mostly involved seriemas that were either far away, not fully within the frame, or recorded with the non-stop cameras, which appeared to produce lower-videoquality footage.

**Figure 3.**
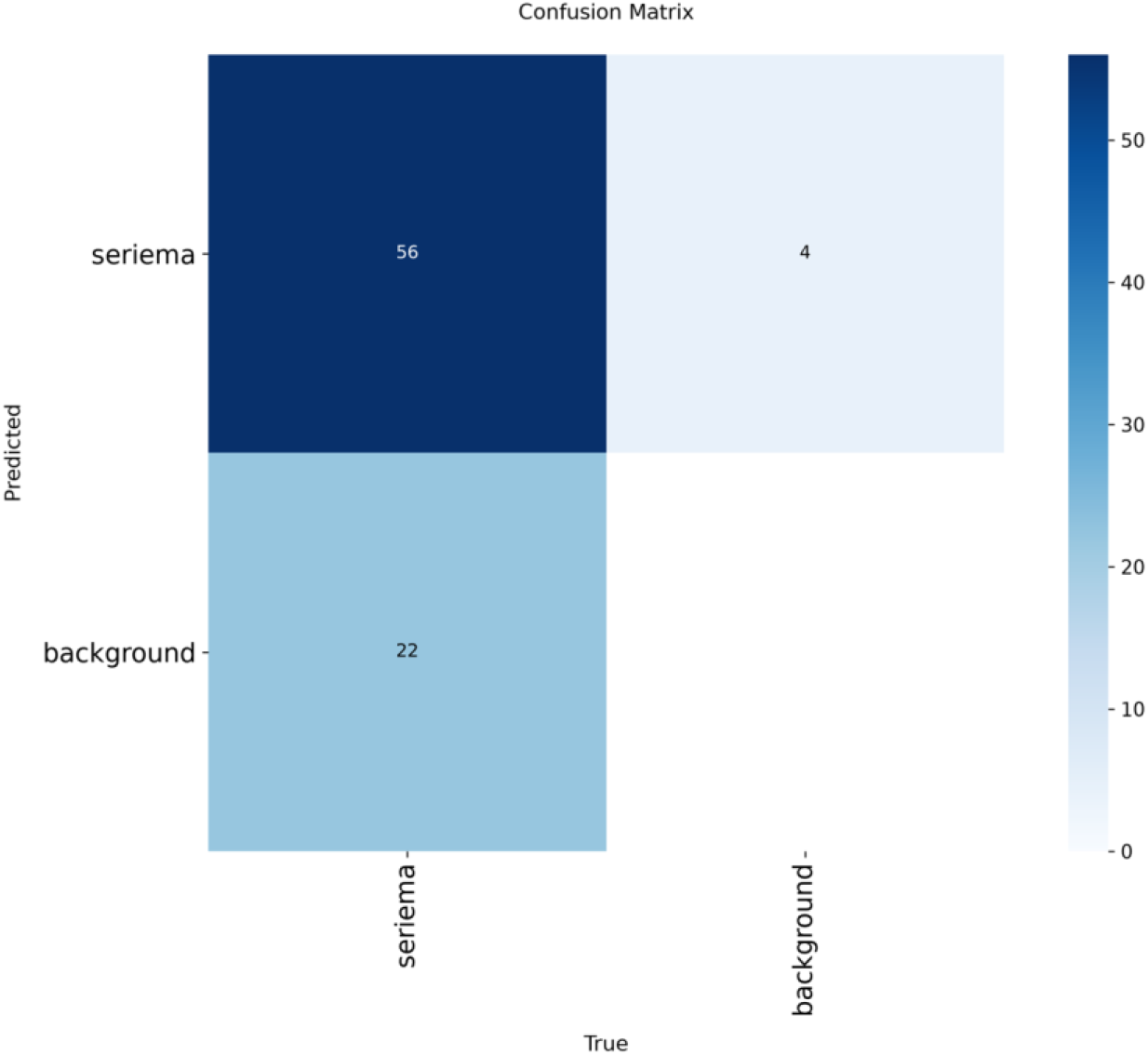
Confusion matrix at a confidence threshold of 25% showing model predictions for the categories seriema and background in the test set, including true positives, false positives, true negatives and false negatives.

The model was subsequently applied to four non-stop camera videos to assess processing speed. At a stride of 10 frames, 716.8 seconds of footage was processed in 156.21 seconds — approximately 4.6 times faster than real time. The model also correctly classified all four videos with respect to the presence or absence of seriemas.

## Discussion

This study provides a workflow for developing a YOLO26-based animal detection model, illustrated here through the detection of red-legged seriemas. A total of 480 images were annotated, of which 288 were used for training, 96 for validation and 96 for testing. The trained model performed relatively well on the validation set, with a precision of 0.934 and a recall of 0.786. On the test set, these values decreased slightly but remained high, with a precision of 0.865 and a recall of 0.740.

Precision reflects a model’s ability to correctly identify the objects of interest — in this case, seriemas. In the test set, when the model flagged a seriema, it was correct approximately 87% of the time. Recall, on the other hand, reflects how many of the objects of interest that were actually present the model managed to detect; here, the model identified 74% of all seriemas present in the test set. For both metrics, higher values indicate better model performance. Because this model is intended to streamline video selection from non-stop recording cameras, missing videos that contain seriemas is considered a high cost. A high recall is therefore particularly valuable in this context.

This model could be further challenged with the usage of a larger dataset, containing more images of seriemas at greater distances and more images from the non-stop recording cameras, as the model seemed to perform less well on these types of images. Additional footage of other animals present in the area might also help to reduce the number of false positives. Nevertheless, for a first attempt using a relatively small dataset, these results are promising and are encouraged to be replicated and scaled up.

A few practical considerations are worth noting for researchers looking to get the best performance from a YOLO26 model: (i) GPUs generally offer faster processing than CPUs, so using a built-in or external GPU, where available, is likely to improve training and inference speed (Han et al., 2020); and (ii) processing time can also be reduced by terminating analysis of a given video once a sufficient number of frames have been labelled as containing a seriema.

Overall, this study presents a case study for producing an animal detection model that can be used to streamline video processing, illustrated here through the detection of red-legged seriemas. Even though the model was built on a relatively small dataset, its performance was considered good, and a usable detection model was successfully created. More importantly, the workflow presented here is not specific to seriemas and can be adapted by other researchers to build similar detection models for other wild species. In doing so, this study aims to lower the barrier for ecologists seeking to apply object detection models.

## Data and code availability

The full code used for the development of this seriema-detection model, including the scripts for frame extraction, annotation conversion and model training and application, is available in the associated GitHub repository: https://github.com/lotjes2002-hue/Seriema-detection-model.git. Additionally, the dataset used for model development and model testing can be found on Zenodo with the following DIO: https://doi.org/10.5281/zenodo.21531067

